# Functional and structural characterization of H105Y mutation in MtrR protein of *Neisseria gonorrhoeae*

**DOI:** 10.1101/2020.07.07.192807

**Authors:** Divya Sachdev, Indu Kumari, Madhu Chopra, Laishram R. Singh, Daman Saluja

## Abstract

MtrR is a negative regulator of MtrCDE efflux pump. Various N-Terminal and C-Terminal mutations have been reported in different multidrug resistant clinical isolates of *Neisseria gonorrhoeae* across the world. Mutations in N-terminal region of MtrR, are known to abrogate its binding with the promoter thereby providing an evidence that this region encodes for DNA binding domain. In contrast, mechanism of action of mutations in C-terminal, known to play a role in protein dimerization and has site for ligand binding, is left unexplored. In the present study, using *in silico* approach, we observed that H105Y mutation affects the conformation of the protein and its binding with penicillin. Purified, recombinant wild type and mutant MtrR were compared for their binding to promoter region and various antibiotics. Fluorescence spectroscopy, CD spectroscopy and dynamic light scattering assay with wild type and mutant MtrR suggested decreased binding of H105Y MtrR with its promoter without affecting protein dimerization, but due to altered conformation of mutant dimer. Our *in silico* results also suggest altered conformation of the mutant dimer protein leading to difference in the posture of homodimer formed and hence altered binding with DNA. Mutant protein also showed stronger binding with various antibiotics: penicillin, ceftriaxone and ofloxacin. Binding of drugs also leads to altered conformation of the protein which may lead to its decreased binding with the promoter DNA.

**Importance:** Mutations in MtrR, a transcriptional repressor of MtrCDE efflux pump have been reported in various multi-drug resistant *Neisseria gonorrhoeae* across the world. We identified that a C-terminal mutation H105Y, outside the DNA binding domain of MtrR, decreases the binding of MtrR with its promoter. We identified that the mutation alters the structure of the dimer, as well as enhances the antibiotic binding. We envisage that the altered structure of MtrR affects its DNA binding thereby increasing efflux of antibiotics and increased resistance. These findings could be used as a guide to design novel drugs that should either not bind to MtrR or could surpass the conformational change and decreased DNA binding.

## Introduction

*Neisseria gonorrhoea*, one of the major sexually transmitted disease causing pathogen, has developed resistance even to the last-line of antibiotics including cefixime, ceftriaxone, and azithromycin-ceftriaxone dual therapy, making it a superbug (1-3). Overexpression of efflux pumps is one of the most important mechanisms employed by the pathogens to develop resistance against various hydrophobic antibiotics. MtrCDE efflux pump plays a protective role by providing resistance to fecal lipids in rectal isolates of *Neisseria gonorrhoeae as well as* expels the hydrophobic drugs from the organism (4-6). The expression of the pump is negatively regulated by MtrR, a 23KDa protein encoded by *mtrR* gene located 250 bp upstream of *mtrC* start site (4, 7, 8), which binds to the pseudo-direct promoter region of MtrCDE pump as two homodimers (9, 10). MtrR was also shown to regulate the expression of various other genes including down-regulating the genes encoding ATP-binding cassette (ABC) transporters. The ABC transporter is predicted to play role in transport of cysteine, an amino acid required for protein and glutathione synthesis and serves as a source of sulfur for various biomolecules such as biotin, coenzyme A, and lipoic acid (11-14). Cysteine and glutathione derivatives help *Neisseria* to combat oxidative stress during infection and thus understanding the mechanism of regulation of genes for cysteine biosynthesis pathway or its uptake by *Neisseria* could shed light on the development of next-generation antimicrobials against the pathogen (15).

MtrR is a member of the TetR family and bears a high degree of structural and functional homology with other members especially with AcrR of *E. coli* and QacR of *Staphylococcus aureus*. Like other family members, MtrR has N-terminal Helix-turn-helix DNA binding domain (amino acid 32-52) while the C-terminal is apparently responsible for dimerization which is a prerequisite for DNA-binding and ligand binding (4, 6, 10, 16-18). It was recently shown that C-terminal of MtrR binds with bile salts, which leads to its reduced binding with the promoter DNA (6). Unlike other family members, MtrR has an additional stretch of 10 amino acids out of which five are basic, present at its N-terminal, before the start of DNA binding site. The stretch forms the helix 1 of MtrR, which resides perpendicular to helix 3 (recognition helix), and is predicted to be involved in electrostatic interaction with the promoter DNA (6, 9).

Mutations both in promoter region (A/T deletion or TT insertion in the 13 bp invert repeat) and in structural region of MtrR (G45D, A39T) have been shown to abrogate binding of MtrR with the promoter of MtrCDE efflux pump and enhance the expression of downstream genes (8, 10, 19-21). Numerous drug-resistant clinical isolates with deletions in the promoter region, point mutations G45D or A39T DNA in N-terminal and H105Y in C-terminal and truncated protein are well reported across the world (22-27) (20)). The role of A/T deletion and G45D in reducing/abrogating MtrR-DNA binding is well understood (7, 8, 10), but the molecular mechanism of H105Y is poorly explored.

In the present study, we have characterized the H105Y mutation with respect to the structure and function of MtrR protein. We hypothesise that this mutation may affect the conformation, dimer formation, and thus the binding of the MtrR with its promoter DNA. Various biophysical assays including fluorescent spectroscopy, CD spectroscopy assay, and dynamic light scattering were used to understand the effect of the mutation on its structure and function. We also studied the effect of binding of various antibiotics on the structure of wild type (WT) and mutant MtrR.

## Methods

### Cloning, expression and purification of His-MtrR fusion proteins

Full-length *mtrR* was PCR amplified with specific primers BamH1 mtrF (5’-<u>GGATCC</u>GCCCTCGTCAAAC-3’) and BstB1 mtrR (5’-TTGTTTGT<u>TTCGAA</u>CGGCA-3’) using gDNA of WT *N. gonorrhoeae*, FA19 strain, (a kind gift by Dr. Fred Sparling, University of North Carolina, USA) (28) and gDNA isolated (as described previously) from an antibiotic resistant clinical isolate with the H105Y mutation (26). Amplification was performed using Taq DNA polymerase in thermal cycler for 36 cycles with the following parameters: 95°C for 30 sec, 60°C for 30 sec, 72°C for 30 sec. The desired PCR products were purified, digested with BamH1 and BstB1 and ligated in pTrc HisA vector (Invitrogen) digested with the same enzymes. The recombinant clone was selected and confirmed by sequencing. Both WT and mutant MtrR were overexpressed in BL21 cells at 30°C for 4h using 0.8 mM IPTG and purified using Ni-NTA affinity chromatography. Briefly, cell pellets were suspended in lysis buffer [20 mM sodium phosphate pH 8; 300 mM NaCl, 10% glycerol, 1 mM Tris (2-carboxyethyl) phosphine hydrochloride (TCEP)], treated with lysozyme (100µg/ml) for 30 minutes at 25°C and sonicated. Supernatant was loaded onto a Ni^2+^ - nitrilotriacetic acid column and hexa-histidine tagged MtrR was purified (>95%) using 300 mM imidazole in lysis buffer. The purified proteins were dialyzed against 20 mM sodium phosphate pH 8, 10% glycerol, 1 mM TCEP overnight to remove the salt. Fractions were analysed using SDS PAGE and Western Blot analysis.

### Protein Cross-linking

Cross-linking reactions were performed in 1X cross-linking buffer (50 mM Tris-Cl pH 8, 100 mM NaCl) using 16 µg of freshly purified MtrR protein (WT or H105Y mutant). Reactions were incubated at 25°C for 20 min. The dimer formed was stabilized by the addition of 1% formaldehyde and incubating the mix at 25°C for 30 min. Reactions were stopped by the addition of 2X SDS loading dye (without β-Mercaptoethanol) and samples were loaded on SDS PAGE. Coomassie staining was performed to analyze the difference in dimer formation.

### Steady-state Fluorescence Measurement

Fluorescence spectra were measured to study the binding of WT and mutant H105Y MtrR protein with DNA and various drugs, using a PerkinElmer LS 55 Spectrofluorimeter, with a quartz cell at 25°C. For all experiments we used protein concentration 0.5 µM – 1 µM, an excitation wavelength of 280 nm and emission spectra was recorded between 290 nm and 420 nm, with both excitation and emission slits set at 5 nm. All spectra obtained were subtracted for the contributions of buffers and necessary controls. Both proteins were incubated for 30 minutes at 25°C with increasing concentration of MtrR promoter DNA (0.1 µM to 5 µM) and ABC promoter DNA (10 µM to 100 µM) in binding buffer (10 mM Tris-Cl pH 8, 100mM NaCl, 1mM DTT), filtered through 0.2 µm sterile filters (Nunc, USA). The dissociation constant (Kd value) was calculated using origin 7 software. Similarly binding of proteins with various drugs was also studied. Proteins were incubated at 37^°^C for 2 hrs with different concentration of taurodeoxycholate (TDCA, a bile salt 2 µM - 50 µM), penicillin (0 µM - 50 mM), amoxicillin (2 µM - 200 µM), ceftriaxone (2 µM - 100 µM) and ofloxacin (2 µM – 100 µM), prepared in syringe filtered 20mM phosphate buffer containing 100 mM NaCl. Effect of the binding of 1-anilino-8-naphthalene sulfonate, ANS (8 µM) on the structure of proteins was also studied.

### Circular Dichroism Spectroscopy

The secondary structure of purified WT MtrR and H105Y mutant MtrR was determined by CD-spectroscopy using Jasco J-810 Spectropolarimeter (163-900 nm RANGE) equipped with a Peltier thermoelectric type temperature control system and flow-through HPLC cell. The measurement was carried out using “far-UV” spectral region (200-240 nm), with cells of 0.1 cm path length at 25°C. Measurements were made at least three times with 3 accumulations of each spectrum. All spectra obtained were subtracted for the contributions of buffers and necessary blanks. The effect of salt, temperature, and binding of various antibiotics on the structure of the purified protein was analyzed. For each experiment, we used 2 µM of purified proteins. To study the effect of salt, proteins were prepared in 20 mM of phosphate buffer (pH 8.0) and incubated with increasing concentrations of NaCl (50 mM to 300 mM) at 4°C and 25°C for 4 hrs and overnight. The optimum salt concentration for maximum stability of protein was identified and used for all subsequent experiments. Thermal stability of WT and H105Y mutant MtrR was also studied by progressively increasing the temperature of the cell from 20°C to 85°C. Three scans were measured over the whole temperature range at 222 nm with dynode voltage kept below 800V.

To study the effect of binding of antibiotics on the structure of MtrR (WT and H105Y mutant), purified proteins were incubated with 50 µM of ceftriaxone and ofloxacin for 2 hrs at 37°C. The CD spectrum of protein treated with antibiotics was measured similarly as for untreated protein.

### Dynamic light scattering

To study the effect of the mutation on dimerization, the dynamic light scattering was performed using a Zetasizer Nano-Zs spectrum (Malvern Instruments, Ltd., U. K.). A 12 µL sample was added in the cuvette, and three measurements of each sample were made with 13 scans for each of them.

### Homology modelling of MtrR

The 3D structure of MtrR of *Neisseria* was modelled to predict the effect of mutation on binding to the promoter region. PDB protein BLAST (http://blast.ncbi.nlm.nih.gov/) was run to find out the highly homologous structures. The PDB structures of AcrR protein of *E. coli* (PDB ID: 2QOP) (16) and QacR of *Staphylococcus aureus*, (PDB ID: 1JTO) (29) were retrieved from Protein Data Bank (www.rscb.org/pdb) and used as template to model the structure of WT MtrR using Discovery Studio 2.5.5 (Accelrys, Inc., San Diego, CA, USA) Although crystal structure of MtrR is also available, due to the presence of CAPS (present in crystallization buffer), in the ligand pocket, structure was in more open conformation and thus could not be used for protein-DNA binding studies (6). Mutant H105Y was generated using the ‘build mutant’ module of Discovery Studio. The modeled structures were typed with CHARMm force-field and refined by simulating under physiological conditions. Water molecules were added to the modeled structure using the explicit solvent model as it brings the structure to room temperature. Counter ions were added to maintain the electroneutrality of the structure. Both the structures were further relaxed (10,000 steps) till the RMSD was 0.0005 kcal/mol/Å using steepest descent and conjugate gradient. Further molecular dynamics simulations were carried out using a leapfrog algorithm with a time step of 1.0 fs. The molecule was slowly warmed from 50K to 300K over a course of 100 ps. After running the heating-cooling cycle, another cycle of equilibration was run for 120 ps at a constant temperature of 300K followed by a production cycle of 120 psec at 300K and constant pressure (NPT) by weakly coupling the system to a thermal bath. The time constant used to couple the system to the thermal bath was 0.1 psec. The structure with the least energy and lowest RMSD was used. Similar steps were also carried out for the mutant protein. The resultant models were checked by drawing the Ramachandaran plot using PROCHECK v.3.0 (30).

### Docking of MtrR with penicillin

Penicillin G was docked into the WT and mutant MtrR using the cDOCKER module of Discovery Studio (31). Both structures were typed with CHARMm force field. It is well known that ligand binds in the cavity formed by helices 4 - 8 in MtrR as well as other family members, thus putative binding site in MtrR was defined in concordance with this information and based on the shape of the cavities of the receptor protein using Define and edit binding site’ tool of Discovery studio (6, 17, 18, 32). Best Pose out of 10 obtained after docking was selected as with minimum cDOCKER interaction energy, minimum binding energy, and maximum Ludi score, and used to study the interaction of proteins with penicillin by drawing interaction diagram.

### Docking of MtrR with its promoter DNA

Model of WT and mutant MtrR dimer were constructed using the ZDOCK protocol of the ‘Protein modelling module’ of discovery studio (33). Up to 2,000 rigid-body docking sites from each program were then filtered, clustered (15 Å clustering radius) including desolvation and electrostatic energies. Best pose with maximum structural similarity with a dimer of the crystal structure of QacR, AcrR and MtrR, maximum ZDOCK score and minimum ZRANK score was selected and clashes were removed by following steps of RDOCK (34). Each selected ZDOCK is run through 130 steps of Adopted Basis Newton-Raphson (ABNR) energy minimization with CHARMm, using the CHARMm force-field. The best pose of WT and mutant dimer selected were docked with DNA. Before docking, DNA was prepared in discovery studio using ‘build and edit nucleotide’ tool and was energy minimized. Double-stranded DNA was typed with ‘Charmm27, partial charge: momany-Rone’. Residues known to involve in docking were selected in order to filter out the poses not involving these residues. Poses with high ZDOCK score and low ZRANK scores and involving residues previously known to bind were selected and RDOCK protocol was run to remove the clashes if any.

## Results and discussion

Mutations in the mtr operon or in its regulatory protein MtrR results in the increased efflux of drugs and thereby make the cell resistant to the said drug. In our initial analysis with mutation patterns in clinical isolates, we observed twenty-two out of 27 isolates (81.4%) carried mutation either in the promoter region or in the coding region of MtrR protein. The most common mutation observed in the MtrR gene was H105Y, present in 12 out of 27 clinical isolates and was mostly coupled with A/T deletion in the promoter region (11/12 isolates) (26). The mutation was also identified from drug-resistant gonococci from other parts of the world (4, 24, 25, 35, 36). Since H105Y lies outside the DNA binding domain, this observation prompted us to understand the molecular basis of resistance in isolates harboring H105Y mutation. In the present study, we have explored the effect of the mutation in MtrR on its conformation, dimerization ability, drug-binding, and DNA-binding ability.

WT and H105Y mutant MtrR protein were successfully cloned, expressed and purified using Ni-NTA chromatography (Fig. S1). All steps were carried out at 4^°^C. Purified protein was dialyzed, concentration was estimated and used within four-days of purification for various biophysical assays. Simultaneously, we used *in-silico* approach to study the effect of mutation on its structure and binding with DNA and various antibiotics.

### Effect of H105Y mutation on the conformation and dimer formation of MtrR

Intrinsic fluorescence of MtrR was used to study the effect of the mutation on its conformation. A prominent decrease in fluorescence of mutant protein suggested that the change in its conformation may be due to burial of aromatic amino-acid inside the hydrophobic core of the protein (Fig. 1a). Prominent peaks at 222 nm and 208 nm were observed in far-UV CD of the WT and mutant protein. Both WT and mutant MtrR were found to be stable in the presence of 100 mM NaCl. The decrease in the dip of peaks of mutant protein suggested loss in helical content (Fig. 1b). We also studied the effect of mutation on thermal stability of the protein and observed WT has higher stability under the physiological range. More than 91% of WT MtrR was stable even at 40°C whereas only 85% of the mutant retained its conformation. We also observed that loss in helical content was sharper in WT above 60°C (Fig. 1c, Fig. S2). MtrR-ANS binding studies also suggested altered conformation of H105Y mutant protein (Fig. 1d). Further, analyzing the effect of the mutation on dimer formation, cross-linking results suggested no change in the dimerization ability of the mutant as compared to the WT protein (Fig. 2). Our data of dynamic light scattering assays suggested altered hydrodynamic radii of the mutant, with the dimer of the mutant being more compact as compared to that of WT (Fig. 3). It suggests although mutation didn’t disturb/abolish the potential of mutant protein to dimerize, it alters the conformation of the dimer. Since the minor change in the radii of protein could affect its ability to fit in the major groove of DNA, it is possible that compact mutant protein binds feebly with promoter DNA and may dissociate easily.

**Figure 1:**
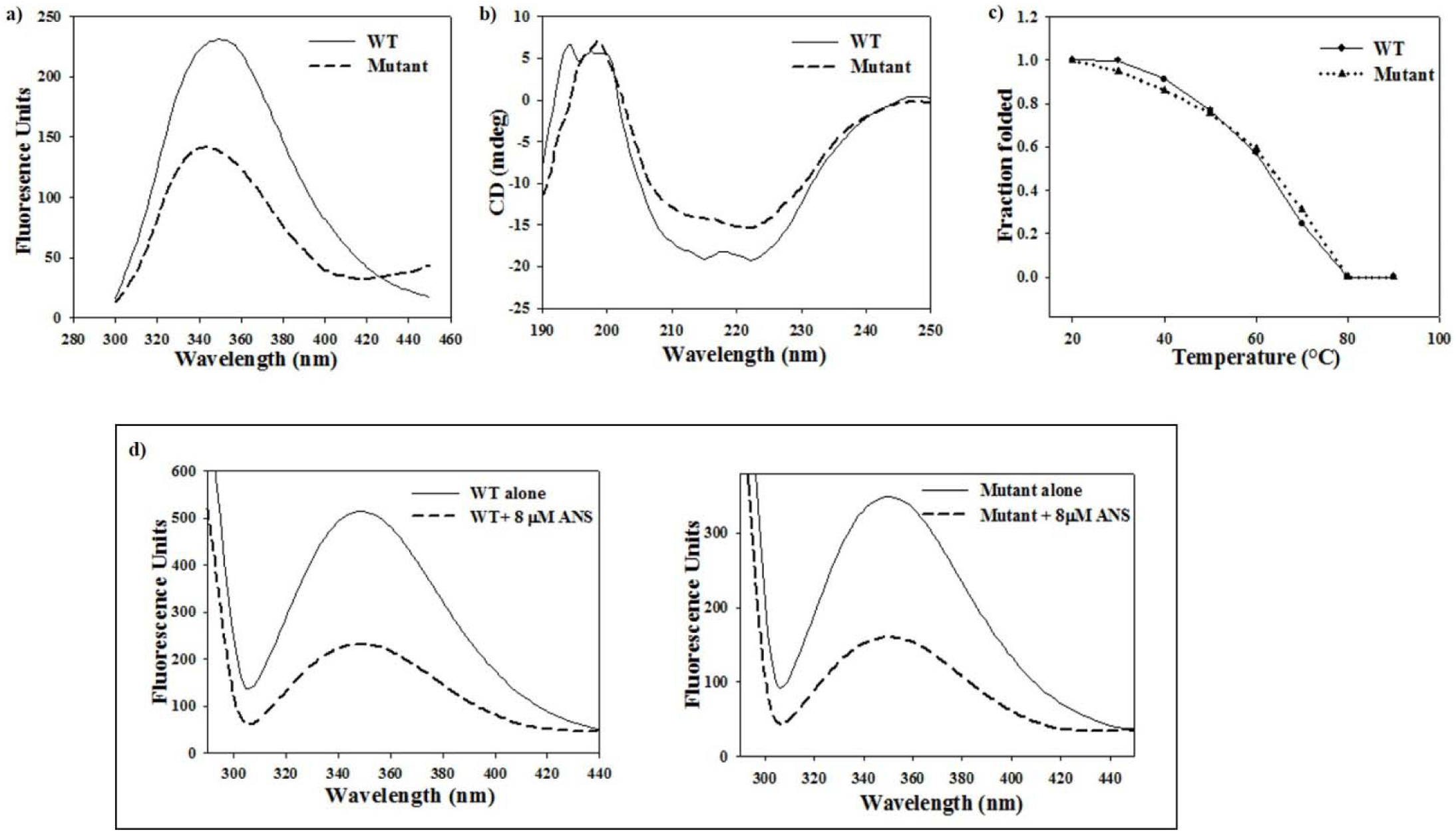
Effect of H105Y mutation on structure of MtrR: a) Fluorescence spectrum of WT and mutant, b) Comparative far-UV CD spectrum of WT and mutant c) Unfolding of WT and mutant on increasing temperature. d) Change in fluorescence on binding of WT (left panel) and mutant MtrR (right panel) with 8 µM of ANS.

**Figure 2:**
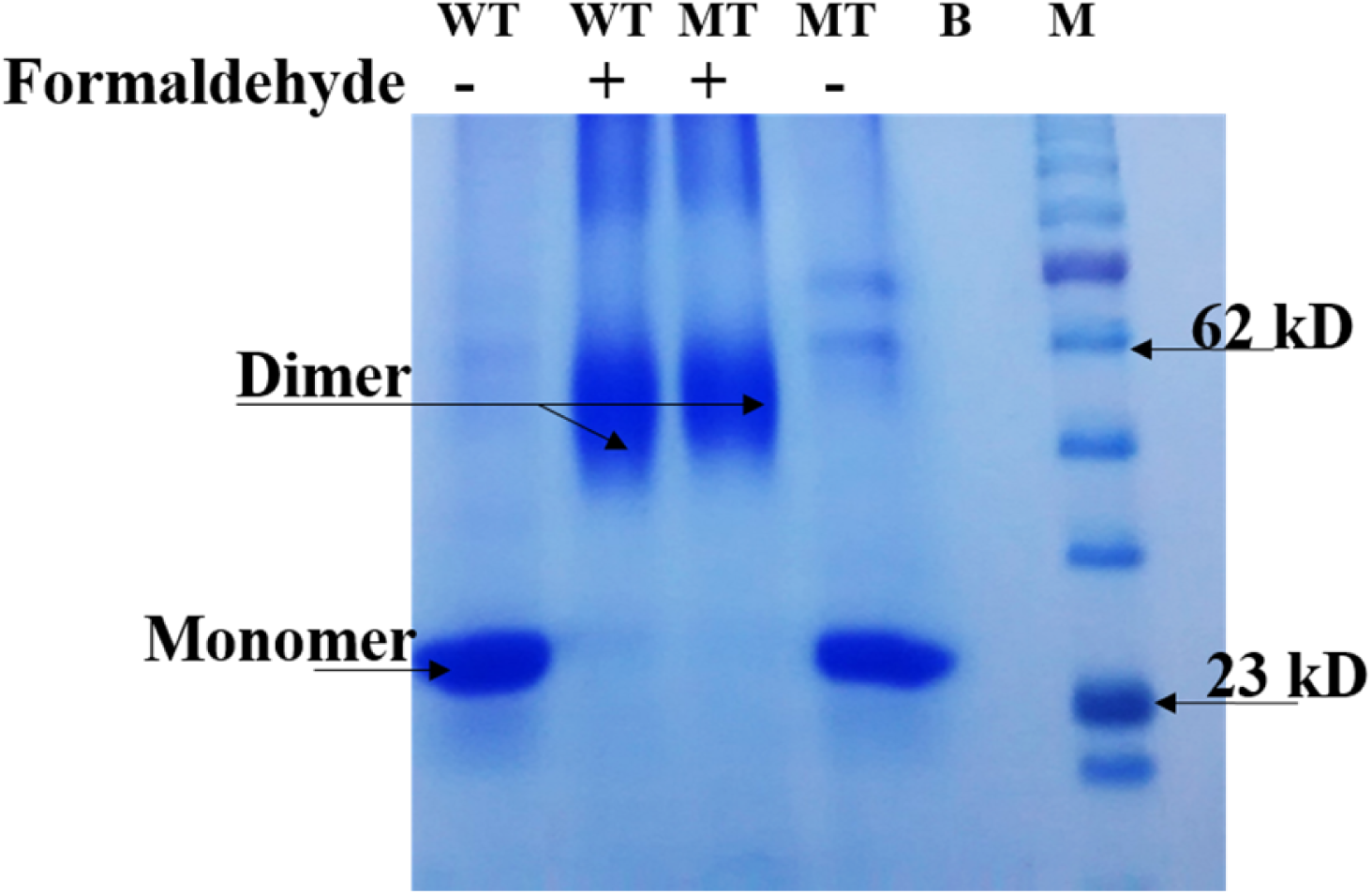
Detection of cross-linked MtrR dimers: Purified wild type and mutant MtrR (16 µg) were cross-linked with 1% formaldehyde separated by SDS PAGE (9%). WT: Wild type, MT: H105Y mutant.

**Figure 3:**
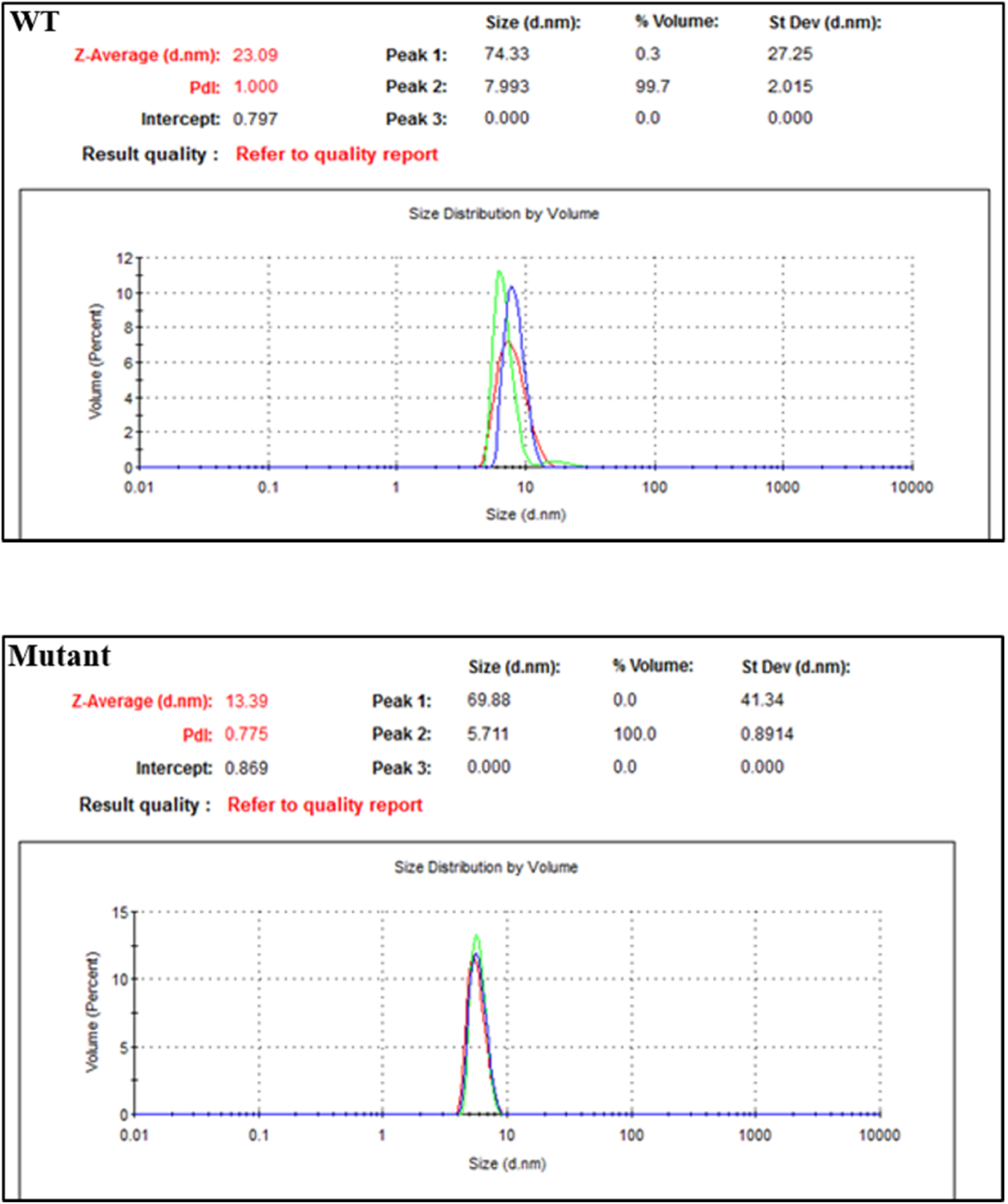
Volume (percent) vs size graph after DLS showing decrease in the hydrodynamic radii of mutant (lower panel) as compare to wild type (upper panel).

Change in conformation was also evident from *in-silico* studies. WT and H105Y mutant of MtrR were build using homology modeling and create mutants module of discovery studio. The final models were validated using the PROCHECK, and more than 90% of residues were found to reside in the allowed region for both the mutant as well as WT MtrR (Fig. S3a and Fig. S3b). Mutant was overlaid with the WT to understand the structural changes induced because of the mutation. The mutation led to a change in the pattern of the hydrogen bond. In WT MtrR, histidine 105 hydrogen bonds with histidine 109 whereas in mutant tyrosine105 forms hydrogen bond with aspargine 102 which resides in the connecting loop between helix 5 and helix 6 (Fig. 4). As a result, helix 5 shifts towards helix 4 (anticlockwise around the central axis) with a concomitant decrease in the volume of the cavity from 356 Å^3^ to 256 Å^3^. These results thus explain the altered secondary structure of mutant H105Y as observed using CD spectrophotometry (Fig. 1b and Fig. S3c) and slight decrease in the size of dimer of the mutant.

**Figure 4:**
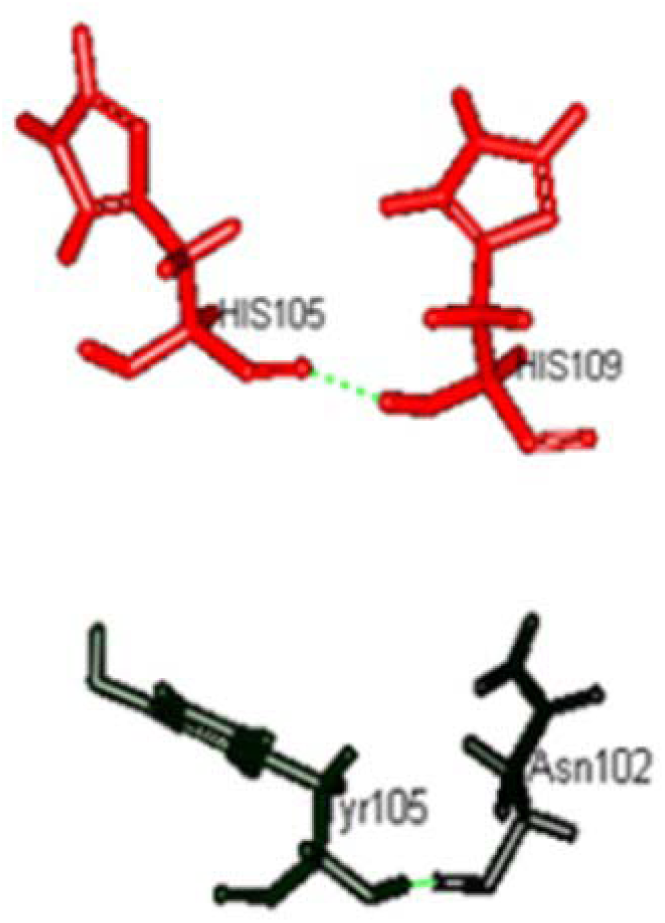
Change in pattern of hydrogen bonding in WT (upper panel) vs H105Y mutant (lower panel) MtrR.

### Mutant MtrR Protein shows decrease in DNA binding affinity and altered binding with various drugs

Wild-type MtrR is well-known to bind to the promoter of MtrCDE efflux pump and has been recently shown to bind with the bile salts. Therefore, it becomes important to study the difference in binding affinity of mutant MtrR for DNA and hydrophobic drugs, with respect to WT. Protein has its intrinsic fluorescence depending on the number of exposed aromatic amino acids. Fluorimetric assays were performed to quantitate the difference in binding affinity of the mutant protein.

Typical hyperbolic spectra were obtained for both WT and H105Y mutant MtrR on titration with increasing concentration of promoter DNA of MtrCDE efflux pump with saturation obtained at higher concentration of DNA for the mutant H105Y than that for WT (Fig. 5, Table 1). The dissociation constant (average of three experiments) was 5.3 µM for WT and 11.5 µM for H105Y mutant MtrR using non-linear fit in origin.

**Table 1:** Normalised fluorescence of WT and H105Y mutant MtrR measures at 340 nm on titration with increasing concentration of fluorescence.

| Wild Type |  | H105Y Mutant |  |
| --- | --- | --- | --- |
| DNA concentration (μM) | Normalised Fluorescence | DNA concentration (μM) | Normalised Fluorescence |
| 0 | 0 | 0 | 0 |
| 1 | 0.2950 | 1 | 0.2200 |
| 2 | 0.4802 | 2 | 0.3300 |
| 3 | 0.5957 | 3 | 0.4400 |
| 4 | 0.6955 | 4 | 0.6600 |
| 5 | 0.8891 | 5 | 0.7600 |
| 7 | 0.9932 | 7 | 0.9800 |
| 8 | 1 | 8 | 1 |

**Figure 5:**
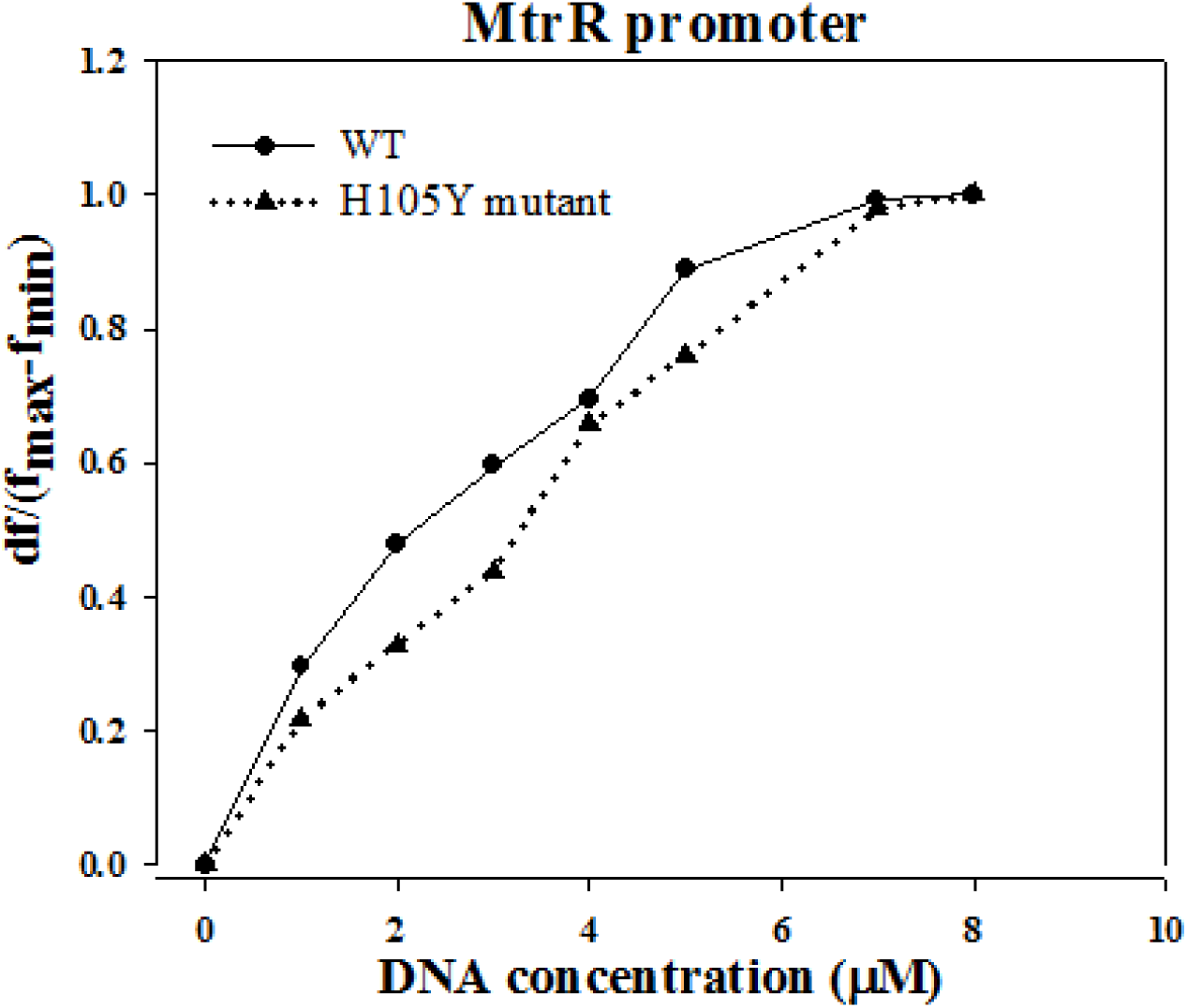
Normalized net fluorescence on of MtrR binding with promoter of MtrCDE efflux pump

It has been recently shown by Beggs et.al that MtrR binds with different bile salts and reduces the binding of MtrR with its promoter (6). We therefore hypothesized that various antibiotics may also bind with MtrR protein and alter its function. To check this, we studied the effect of binding of WT and H105Y mutant MtrR protein with commonly used antibiotics in India to treat gonococcal infection; Amoxicillin, Ceftriaxone, Penicillin, Ofloxacin, and Streptomycin. To speculate the effect of binding of antibiotics on the conformation of the protein, WT and mutant proteins were titrated with increasing concentration of antibiotics. A decrease in fluorescence along with a bathochromic shift (from 340 nm to 360 nm) was observed in the fluorescence of both WT and mutant MtrR when allowed to bind with the increasing concentrations of penicillin. The shift was more pronounced in mutant than in WT MtrR suggesting a pronounced conformational change in mutant as compared to that of WT (Fig. 6a). Similarly, mutant protein has a higher affinity for both Ceftriaxone and Ofloxacin (Fig. 6b and 6c). Neither WT nor mutant MtrR showed binding with Amoxicillin and Streptomycin (data not shown). Loss in the helical structure on incubation with drugs as observed by CD spectroscopy further supports the binding of both WT and MtrR with ceftriaxone and ofloxacin. Loss in the helical structure was more prominent on binding with ceftriaxone (Fig. 7). The higher affinity of mutant with drugs could be because of the altered size of the hydrophobic core which forms the ligand-binding pocket and difference in the binding pattern of drug with the mutant protein.

**Figure 6:**
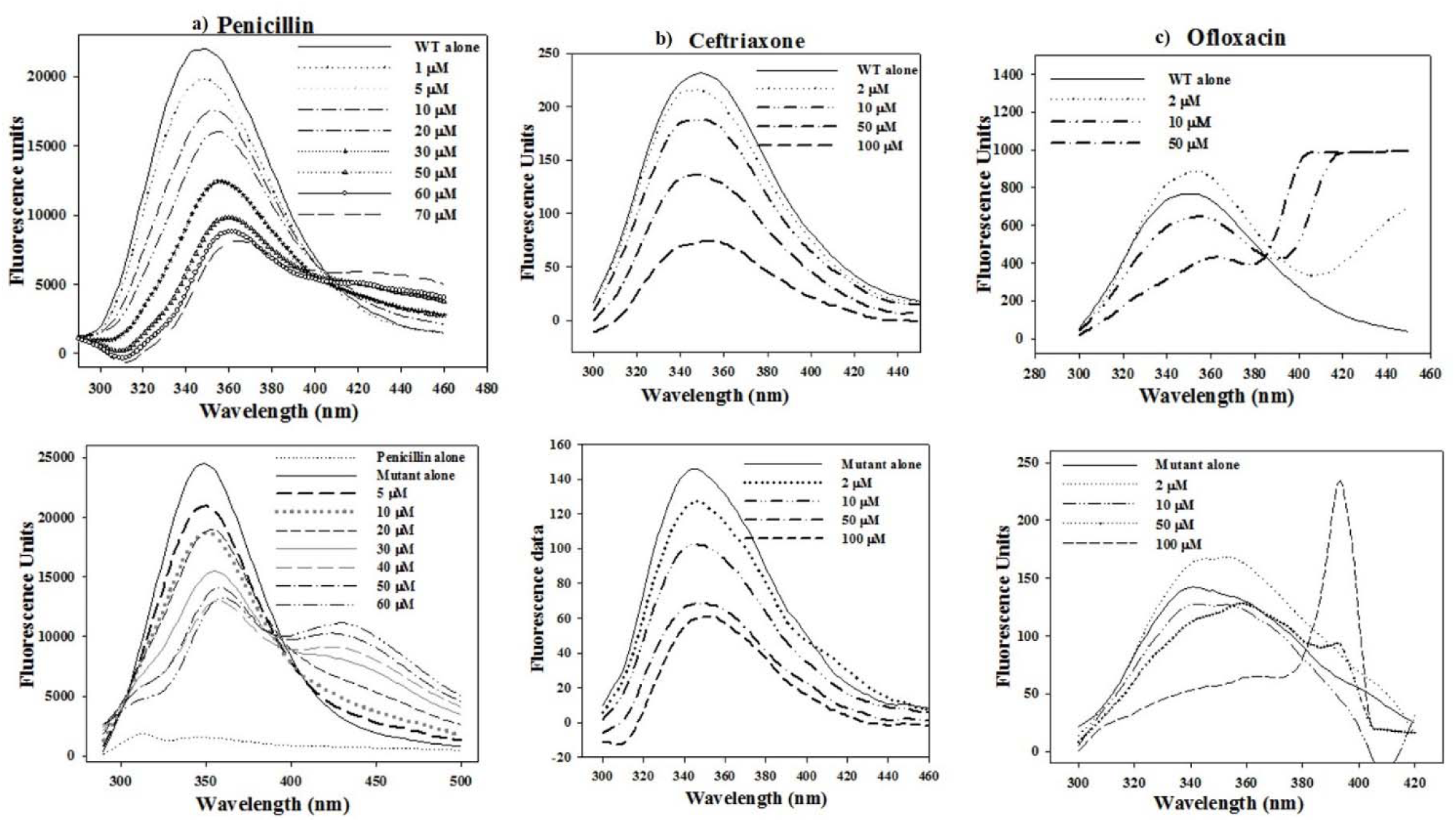
Comparative fluorescence of MtrR on binding with increasing concentration of: a) penicillin, b) Ofloxacin and c) ceftriaxone; Upper panel: WT, Lower Panel: H105Y mutant

**Figure 7:**
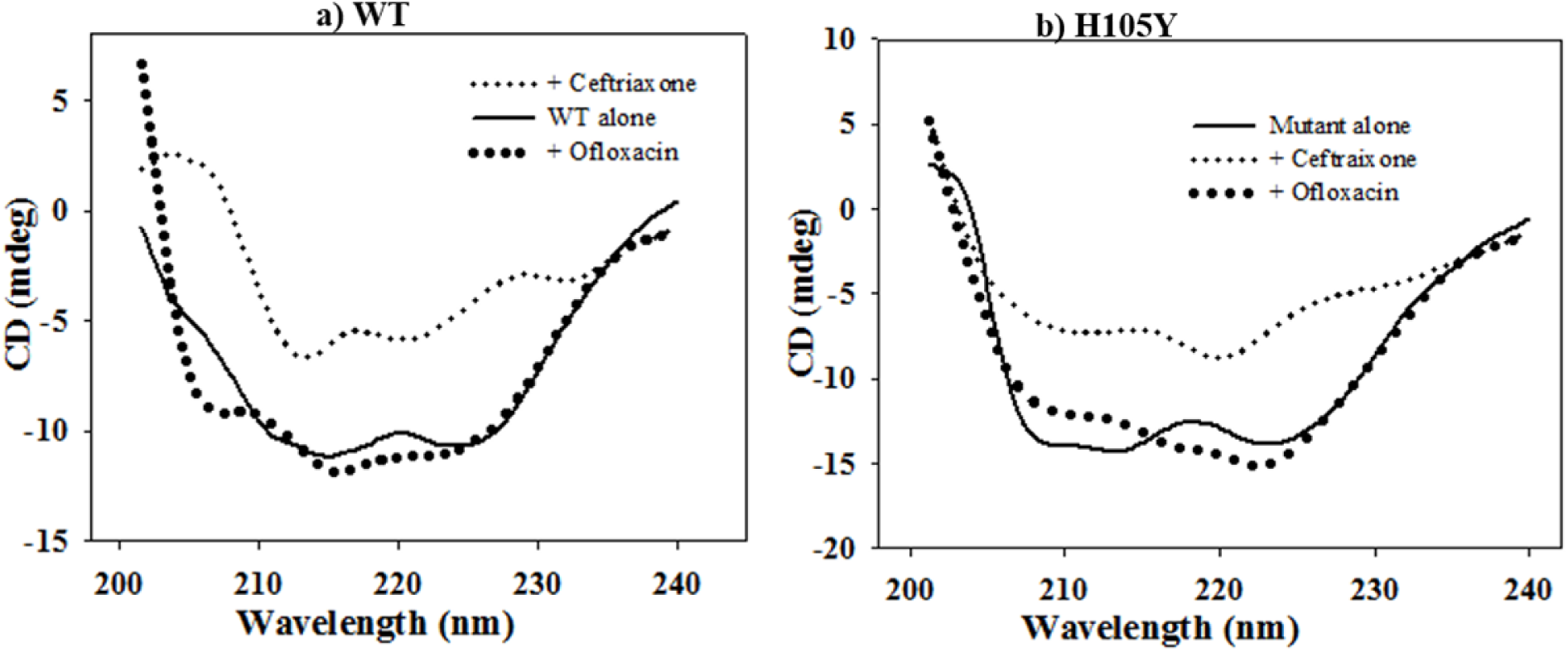
CD spectrum of MtrR on binding with 50 mM of Ofloxacin and ceftriaxone; a) WT, b) mutant

Thus, to understand the molecular basis of difference in binding affinities, we analyzed the binding pattern of penicillin with WT and mutant MtrR using *in-silico* approach. Penicillin was docked to WT and mutant MtrR. Our docking results showed a difference in the pattern of binding of penicillin with WT and H105Y mutant MtrR protein (Table 2a, Fig. 8 and S3d). The key ligand-binding residues (Lys 167 and Cys 66) as found in other functionally similar proteins AcrR, were found within a distance of 5Å of the docked site. Residues Phe 62, Ile 65, Phe 95, Leu 112, Phe 113, Ala 29, Arg 130, His 132, Gln 133, Ile 135 and Trp 136 were found within a distance of 3.5 Å from the ligand which are reported in Table 2a and Fig. 8.

**Table 2a:**
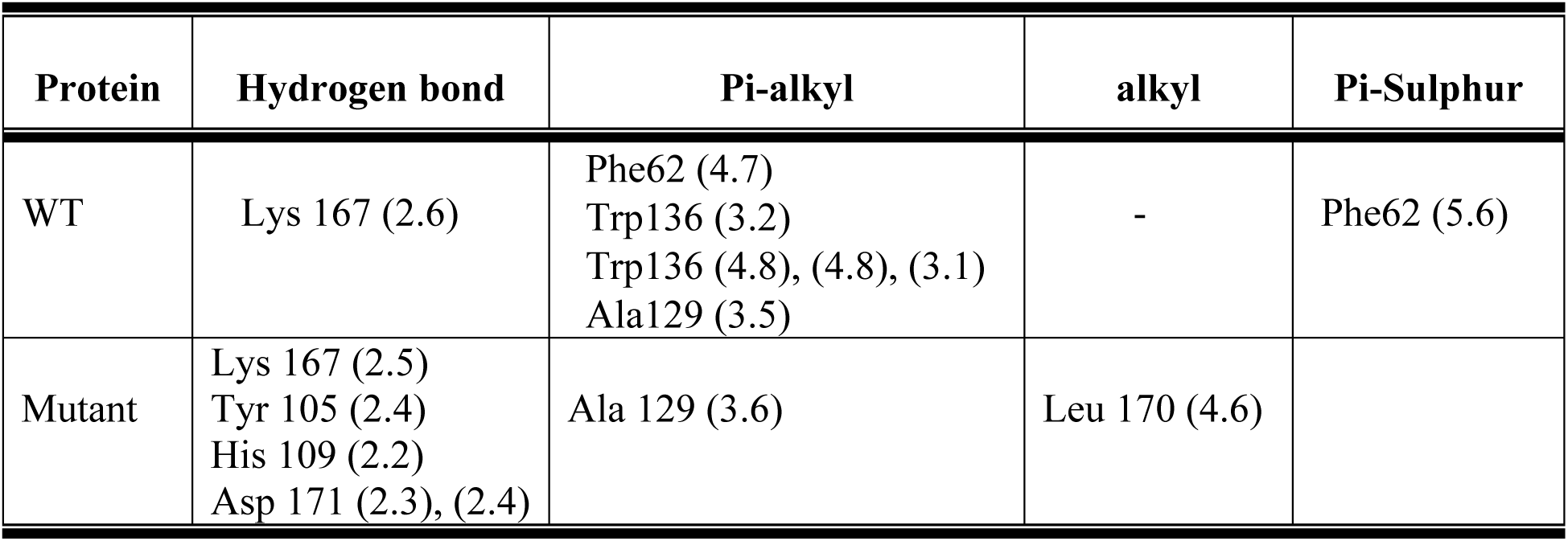
Various iinteractions between amino acids of WT and Mutant MtrR with Penicillin. Amino acid (bond length in Å)

| Protein | Hydrogen bond | Pi-alkyl | alkyl | Pi-Sulphur |
| --- | --- | --- | --- | --- |
| WT | Lys 167 (2.6) | Phe62 (4.7)<br>Trp136 (3.2)<br>Trp136 (4.8), (4.8), (3.1)<br>Ala129 (3.5) | - | Phe62 (5.6) |
| Mutant | Lys 167 (2.5)<br>Tyr 105 (2.4)<br>His 109 (2.2)<br>Asp 171 (2.3), (2.4) | Ala 129 (3.6) | Leu 170 (4.6) |  |

**Figure 8:**
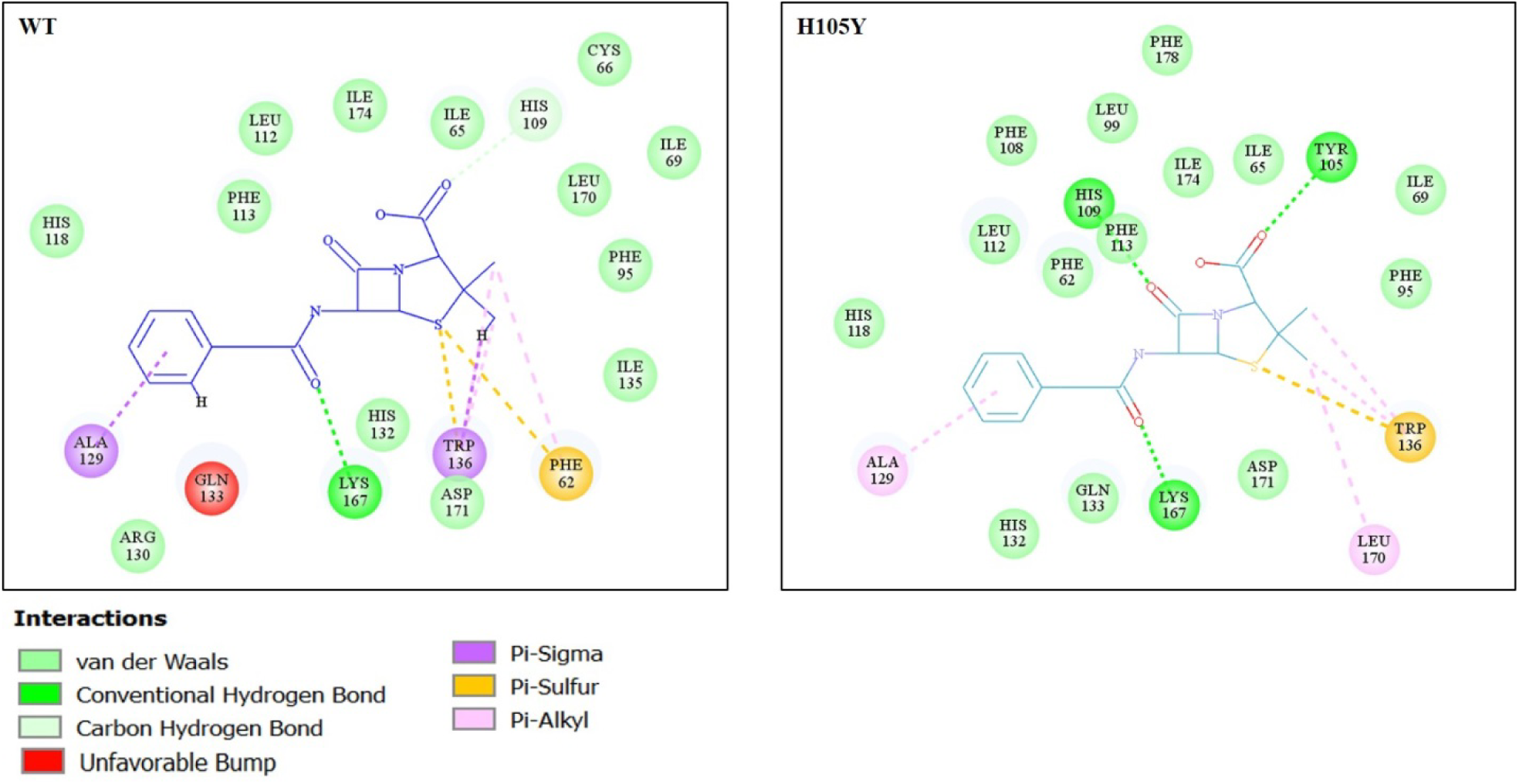
2D diagram showing various interactions of penicillin with amino-acids of WT (left) and H105Y mutant (right) MtrR.

In WT MtrR, ‘=O’ of N-acyl group in penicillin forms one hydrogen bond with Lys 167. The benzyl ring of N-acyl group of penicillin is involved in ‘pi-alkyl’ interaction with Ala 129. Penicillin is also involved in four pi-alkyl interactions with Phe 62 (one) and Trp 136 (three). Another ‘pi-sulphur’ interaction is seen among polar ‘S’ of thiazolidine ring of penicillin and Phe 62. Although Pi-sigma interactions (Pi-alkyl and Pi-sulphur) help penicillin in intercalating in the binding site of the MtrR protein, presence of Gln133 also introduces steric hindrance. The steric hindrance is introduced after in-situ minimization of docked pose suggesting feeble binding of penicillin with WT MtrR. Like in WT MtrR, Lys 167 of mutant H105Y MtrR forms a hydrogen bond with ‘=O’ of N-acyl group in penicillin but with shorter bond length. The benzyl ring of penicillin is also involved in pi-alkyl interaction with Ala 129. Unlike WT, in H105Y mutant MtrR, Tyr 105 (residue of interest) forms an additional conventional hydrogen bonds with penicillin whereas penicillin forms three carbon hydrogen bonds with His 109 (one) and Asp171 (two). Another hydrophobic (alky-alkyl) interaction is observed between Leu170 and penicillin.

The shorter bond length and additional hydrogen bonds in mutant MtrR helps in better intercalating and stronger binding of the penicillin. The formation of a strong hydrogen bond with the substituted tyrosine in the H105Y mutant explains the plausible role of mutant in the binding of penicillin and other antibiotics. Although the mutant has comparable binding energy with that of the WT MtrR, a much higher Ludi score and –cDOCKER energy as compared to that of WT, suggests that the H105Y mutant has a higher affinity for penicillin (Table 2b).

**Table 2b:** Comparison of scoring functions of best pose of WT and mutant MtrR obtained after docking with Penicillin. All energies expressed as (Kcal/mol).

| Protein | -cDock<br>energy | -cDock<br>interaction energy | Ludi 1 | Ludi 2 | Ludi 3 | Binding<br>energy |
| --- | --- | --- | --- | --- | --- | --- |
| Wild type | 20.7 | 36.5 | 350 | 330 | 549 | -150.801 |
| H105Y mutant | 22.68 | 40.62 | 500 | 453 | 570 | -146.5 |

### *In-silico* DNA protein docking to understand protein-DNA interaction

Further, we also studied the molecular basis of the altered binding of the mutant MtrR with the promoter DNA. Since MtrR binds as a dimer with DNA (9), two monomers of WT and H105Y mutant were docked to form dimer using the ZDOCK module. Poses with maximum similarity with QacR dimer, minimum ZRANK score, and maximum ZDOCK score were selected for both the proteins and further refined manually following RDOCK protocol. Dimer formed is stabilized by hydrophobic helix to helix interaction between the helices especially between helix 8 and helix 9 with their counter partner helix8’ and helix 9’ in the second monomer (Fig. S4). Comparing the total energy of the dimer complex, the H105Y complex is found to be a more stable structure Also, mutant dimer has increased center to center distance between G45 of α3 and G45 of α3’ (residue shown to be important in DNA binding) (Table 3). We envisage that change in DNA binding with H105Y mutant could be ascribed to the altered conformation of the dimer. To check this hypothesis, protein-DNA docking was performed using the ZDOCK module. In the absence of crystal structure of WT MtrR complexed with DNA, the best pose was screened using structural similarity with QacR (complexed with its promoter), an another member of the TetR family. It is well established that helix 2 and helix 3 are involved in DNA binding, with helix 3 being the prime region of DNA binding among various members of TetR family (16, 29, 32). Also, Lucas and co-workers have shown using EMSA experiments that mutation in Gly45 amino acid (resides in helix 3) and ADE14, ADE15 and CYT17 of the promoter play important role in-binding (10). Therefore, pose with minimum energy and showing maximum DNA interactions with amino acids residing in helix 3 were selected. Mutant MtrR docked with DNA was superimposed on WT docked with DNA to study change in binding pattern (Fig. S4). The interactions between MtrR (WT and mutant) with its promoter were studied by analyzing the changes in the interaction and conformation of the protein-DNA complex. Several changes in the H-bonding of the protein–DNA complexes were observed as depicted by the differences in the distribution of hydrogen bonds between the interface of macromolecular assembly, i.e., between protein and DNA. The change in pattern of hydrogen bond in WT vs mutant MtrR is tabulated in Table 4.

**Table 3:** Parameters studied to analyze the stability of the dimer complex of WT and mutant MtrR.

| Function studied | WILD TYPE | H105Y mutant |
| --- | --- | --- |
| ZDOCK SCORE | 47 | 9 |
| ZRANK SCORE | -87 | -92 |
| Total energy | -396,122 Kcal/mol | -502,622 Kcal/mol |
| Centre to centre distance | 44.99 | 55.349 |
| Potential energy | 43648 Kcal/mol | 31800 Kcal/mol |

**Table 4:** Comparative analysis of hydrogen bond pattern and various other interactions between WT MtrR DNA complex and H105Y mutant MtrR DNA complex.

| WT |  | MT |  |
| --- | --- | --- | --- |
| Bond Name | Bond length (Å) | Bond Name | Bond length (Å) |
| **Conventional hydrogen bonds** |  |  |  |
| A:THR4:HT2 - S:CYT13:O2 | 2.34478 | A:LYS10:HZ2 - S:CYT13:O3' | 2.6136 |
| A:THR11:HG1 - S:ADE15:O1P | 2.00707 | A:HIS14:HE2 - S:ADE15:O2P | 1.93238 |
| A:THR43:HG1 - S:THY16:O2P | 2.08139 | A:THR43:HN - S:CYT13:O2P | 1.87116 |
| A:HIS50:HE2 - S:ADE15:O5' | 2.05563 | A:THR43:HG1 - S:CYT13:O2P | 1.91446 |
| A:GLN121:HE21-S:GUA24:O1P | 2.62323 | S:ADE14:H61 - A:GLY38:O | 2.65304 |
| A:THR4:OG1-S:GUA12:H21 | 2.0203 | S:ADE15:H61 - A:GLN35:O | 2.52619 |
| A:THR4:HG1 - AS:ADE21:N3 | 2.46589 | A:THR4:HG1 -AS:GUA23:O5' | 2.07358 |
| A:THR4:HT3 - AS:CYT20:O2 | 2.17777 | A:GLU7:HN - AS:GUA23:O1P | 2.78635 |
| A:THR7:HG1 - AS:CYT22:O1P | 1.93515 | A:ARG25:HH22 - AS:CYT12:O5' | 2.0438 |
| A:SER32:HG - AS:CYT12:O2P | 1.88186 | A:ASN34:HD21 - AS:GUA15:O2P | 2.04436 |
| A:ASN34:HD21 -AS:ADE11:O5' | 2.6266 | A:GLN38:HE21 - AS:ADE16:N7 | 2.35357 |
| A:ASN34:HD22-AS:ADE11:O2P | 2.41784 | A:GLN35:HE21 - AS:THY17:O4 | 2.49251 |
| A:TRP49:HE1 - AS:GUA15:N7 | 2.49042 | B:TYR48:HH - S:THY27:O4 | 2.50931 |
| A:ASN53:HD21-AS:ADE13:O1P | 2.14177 | B:LYS54:HZ1 - S:GUA24:O5' | 1.93442 |
| A:LYS54:HZ3 - AS:CYT12:O5' | 2.37941 | B:ARG44:HN - AS:THY4:O2P | 2.25692 |
| B:ARG30:HE - S:THY25:O2P | 2.01399 | B:ALA46:HN - AS:THY3:O2P | 2.36009 |
| B:GLY45:HN - AS:ADE2:O1P | 2.15198 | B:TYR48:OH-AS:ADE5:H61 | 2.78651 |
| **Carbon hydrogen bond** |  |  |  |
| A:ALA8:CA - S:ADE15:O1P | 3.27908 | S:CYT13:H5" - A:GLU7:OE2 | 2.56264 |
| A:THR7:O-S:ADE14:H5' | 2.25221 | S:CYT13:H4' - A:GLU7:OE1 | 2.86905 |
| A:THR4:CA - AS:ADE21:O4' | 3.33946 | S:ADE14:H8 - A:ALA40:O | 2.74515 |
| A:TRP49:CD1 - AS:GUA15:O6 | 3.20507 | S:ADE15:H8 - A:ALA39:O | 2.84874 |
| A:LYS52:O-AS:ADE13:H3' | 2.95669 |  |  |
| A:THR7:OG1-AS:ADE21:H5" | 2.66859 |  |  |
| A:THR7:OG1-AS:ADE21:H4' | 2.32735 |  |  |
| B:ARG33:CD - AS:ADE2:O2P | 3.33666 |  |  |

| Salt bridge |  |  |  |  |
| --- | --- | --- | --- | --- |
|  | A:LYS5:HZ3 - S:ADE14:O1P | 1.88536 | A:LYS10HZ3 - S:ADE14:O1P | 2.00084 |
|  | A:LYS12:HZ1 - S:ADE15:O1P | 2.47676 | A:THR4:HT1 - AS:GUA24:O1P | 2.14342 |
|  | A:LYS12:HZ2 - S:ADE15:O1P | 2.61077 | A:THR4:HT3 - AS:GUA24:O1P | 2.44721 |
|  | A:LYS12:HZ2 - S:ADE15:O2P | 2.61551 | A:ARG25:HH21 - AS:ADE13:O2P | 2.06136 |
|  | A:LYS12:HZ3 - S:ADE15:O2P | 2.34347 | A:LYS26:HZ1 - AS:CYT14:O2P | 2.03731 |
|  | A:ARG44:HH21-AS:ADE11:O2P | 3.20384 | A:LYS26:HZ2 - AS:CYT14:O1P | 2.00319 |
|  | A:ARG44:HH22 - AS:ADE11:O2P | 3.11053 | A:ARG30:HH12 - AS:ADE13:O1P | 2.0107 |
|  | A:LYS52:HZ1 - AS:CYT14:O2P | 2.1099 | A:ARG30:HH22 - AS:ADE13:O1P | 1.97793 |
|  | A:LYS54:HZ3 - AS:CYT12:O1P | 2.72644 | B:LYS52:HZ1 - S:ADE26:O1P | 2.11448 |
|  | B:ARG30:HH12 - S:GUA24:O2P | 2.17863 | B:LYS52:HZ3 - S:ADE26:O2P | 2.02116 |
|  | B:ARG44:HH11 - AS:ADE2:O2P | 1.99204 | B:LYS52:HZ3 - S:THY25:O2P | 1.9468 |
|  |  |  | B:ARG44:HH12 - AS:ADE5:O2P | 2.80748 |
|  |  |  | B:ARG44:HH22 - AS:ADE5:O2P | 2.08812 |
| Electrostatic attractive charges |  |  |  |  |
|  | A:LYS12:NZ - S:ADE14:O1P | 5.40691 | A:LYS10HZ3 - S:ADE14:O1P | 2.00084 |
|  | A:LYS54:NZ - AS:ADE13:O1P | 4.6934 | A:THR4:HT1 - AS:GUA24:O1P | 2.14342 |
|  | B:ARG30:NH2 - S:GUA24:O1P | 4.1477 | A:THR4:HT3 - AS:GUA24:O1P | 2.44721 |
|  | B:ARG30:NH2 - S:THY25:O2P | 3.26516 | A:ARG25:HH21 - AS:ADE13:O2P | 2.06136 |
|  |  |  | A:LYS26:HZ1 - AS:CYT14:O2P | 2.03731 |
|  |  |  | A:LYS26:HZ2 - AS:CYT14:O1P | 2.00319 |
|  |  |  | A:ARG30:HH12 - AS:ADE13:O1P | 2.0107 |
|  |  |  | A:ARG30:HH22 - AS:ADE13:O1P | 1.97793 |
|  |  |  | B:LYS52:HZ1 - S:ADE26:O1P | 2.11448 |
|  |  |  | B:LYS52:HZ3 - S:ADE26:O2P | 2.02116 |
|  |  |  | B:LYS52:HZ3 - S:THY25:O2P | 1.9468 |
|  |  |  | B:ARG44:HH12 - AS:ADE5:O2P | 2.80748 |
|  |  |  | B:ARG44:HH22 - AS:ADE5:O2P | 2.08812 |
|  |  |  | B:ARG44:HH12 - AS:ADE5:O2P | 2.80748 |
|  |  |  | B:ARG44:HH22 - AS:ADE5:O2P | 2.08812 |
| Hydrophobic |  |  |  |  |
|  | A:HIS50-S:ADE15:O1P<br>(Pi-Anion) | 4.17383 | S:ADE14:O1P - A:HIS14<br>(Pi-Anion) | 4.71866 |
|  | A:TRP49-S:ADE15:H61<br>(Pi-Donor Hydrogen Bond) | 2.86005 | A:GLY41:CA - SADE14<br>(Pi – Sigma) | 3.70296 |
|  | A:TRP49 - S:THY16<br>(Pi-Pi T-shaped) | 5.90557 | S:ADE14:H5" - A:HIS14<br>(Pi – Sigma) | 2.65916 |
|  | A:TRP49 - S:ADE15<br>(Pi-Pi T-shaped) | 5.77984 |  |  |
|  | A:TRP49-S:ADE14<br>(Pi-Pi T-shaped) | 5.09389 |  |  |

Considering the bonds with ADE14 of the sense strand, mutating which has been shown to decrease the DNA-MtrR binding (10), its phosphate (O1P) formed a salt bridge and was involved in electrostatic attraction with the positively charged lysines (Lys5 and Lys7 respectively) in the N-terminal of WT protein. ADE14 was involved in carbon hydrogen bond with Thr7 and showed pi-pi T-shaped hydrophobic interaction with Trp49 of WT MtrR. Unlike WT, ADE14 is involved in one salt bridge (Lys10), one carbon-hydrogen bond (Ala40) and three pi-orbital interactions (Table 4). Homologue amino acid of Trp49 of MtrR among various other TetR family members including Trp43 in TetR, Trp50 in AcrR and Tyr41 in QacR are shown to play a role in DNA binding (16, 29, 32) as well as positively charged residues at extreme N-terminal has been predicted analogous to positively charged residues of SimR and AmtR, two other members of TetR family shown to play important role in DNA binding (37, 38). Thus, difference in bonding pattern of ADE14 and vital amino-acids in mutant MtrR with respect to WT protein suggests one plausible reason for reduced affinity of the mutant for DNA. Also, ADE15 of the sense strand, mutating which enhanced the binding (10), formed 2 hydrogen bonds and 4 salt bridges with the WT protein whereas the H105Y mutant was able to form only one hydrogen bond with ADE15.

Similarly, in WT MtrR, one of the critical amino-acids in DNA binding, Gly45 of distal monomer forms a hydrogen bond with phosphate of ADE2 of antisense strand whereas this residue is not involved in hydrogen bond formation in the mutant MtrR. Mutating Gly45 to Asp (G45D) was shown to abrogate the binding of MtrR with its promoter DNA (Lucas) as well as was shown important for tight docking of QacR (Gly37) with its promoter, suggesting role of Gly45 in stronger binding of WT MtrR (29).

Also, Lys36 in QacR and Arg28 in TetR were shown to play imperative role in the binding of repressors with their promoter DNA (29,39). Our *in-silico* results also showed that Asn34 (Arg 28 in TetR) of proximal monomer forms two hydrogen bonds with the DNA whereas in H015Y mutant it is involved in the formation of only one hydrogen bond. Arg44 (Lys36 in QacR) of both the monomers of MtrR is involved in the formation of three salt bridges with promoter DNA in WT MtrR whereas in mutant MtrR, only distal monomer is involved in the formation of two salt bridges and one hydrogen bond. The amino-acid is also involved in various electrostatic interactions with DNA among mutant protein. Involvement of Arg44 of both the monomers in WT may add to its stronger binding with the promoter DNA.

Various other amino-acids including Thr25, Ser34, Ser35, Asn38, Thr40, Th441, His42 and Lys46 in QacrR and Tyr49, Trp50, His51 and Lys55 in AcrR are also shown to play an important role in the binding of repressors to their promoter (16, 17, 29, 32). In our study, we observed variation in hydrogen bond pattern of the corresponding amino-acids in MtrR among WT and mutant, and are thus expected to play a significant role in altered affinity of mutant MtrR for the promoter DNA.

WT MtrR forms 12 hydrogen bonds with phosphates of nucleic acid bases which play major role in stabilization of DNA-protein interactions whereas mutant forms only 10 hydrogen-phosphate bonds again suggesting weaker binding of the mutant. WT MtrR forms higher number of hydrogen bonds (conventional and carbon hydrogen) and pi-bonds whereas mutant bond pattern is dominated by salt bridges and electrostatic attractive bonds. This altered alignment of mutant protein on DNA may be responsible for decreased affinity with a promoter which would result in altered gene expression of the operon.

## Conclusion

Fluorescence assay, DLS assay and CD spectroscopy data suggested altered conformation and decreased binding of H105Y MtrR with its promoter DNA. The results are also supported by *in-silico* data. The increased center to center distance between the DNA recognition helices of two monomers (i.e α3 and α3’) of the mutant MtrR changes the alignment of protein on DNA. We envisage that the monomer is unable to fit in the major groove of DNA and thus contact different DNA bases. The change in the hydrogen bond pattern formed between critical amino acids in H105Y mutant MtrR and DNA bases may be sufficient to explain this decreased protein-DNA binding of the H105Y mutant of MtrR observed using fluorescence spectroscopy. Also, mutant protein also showed stronger binding with various antibiotics including penicillin, ceftriaxone and ofloxacin. It has recently been shown that binding of bile salts leads to altered conformation of homodimer and thus reducing its binding with promoter DNA (6). Our CD spectroscopy results showed that binding of drugs leads to altered conformation of the protein which may result in decreased binding with the promoter DNA. Thus, even a single amino acid change, outside the DNA binding domain or drug interacting/binding domain, is sufficient to cause a conformational change in the protein which results in altered stability of the protein-DNA complex. An obvious consequence of altered DNA binding will be increased expression of the proteins of the efflux pump leading to antibiotic resistance.

## Acknowledgements

Funding from University Grant Commission, India (UGC, Project Grant No. 36-254/2008 & UGC-SAP II project to DS) and DBT bioinformatics facility (DBT-BIF) of ACBR (Project Grant No. BT/BI/25/038/2012) is acknowledged. D. Sachdev and I. Kumari are grateful to Indian Council of Medical Research for Senior Research Fellowship. A kind gift of wild type strain (FA19) of Neisseria *gonorrhoeae* by Dr. Fred Sparling, University of North Carolina, USA is greatly acknowledged.

## Supplementary figures

**Figure S1:**
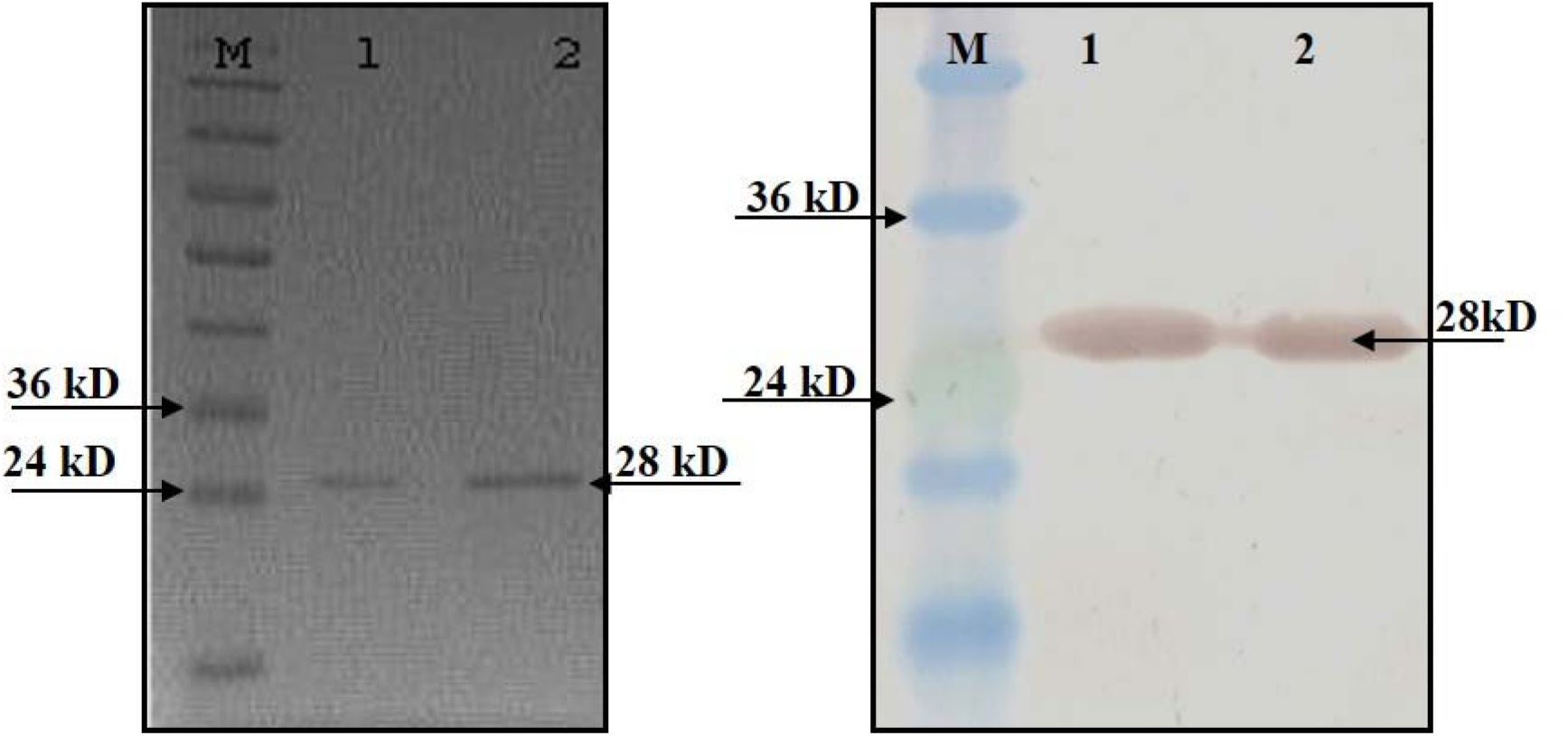
Purification of MtrR using Ni-NTA chromatography: **Left** Commassie stained SDS-PAGE gel of recombinant MtrR (WT and mutant in Lane 1 &2); Right panel: Purified MtrR was confirmed using anti-His antibodies. M: Protein molecular weight marker

**Figure S2:**
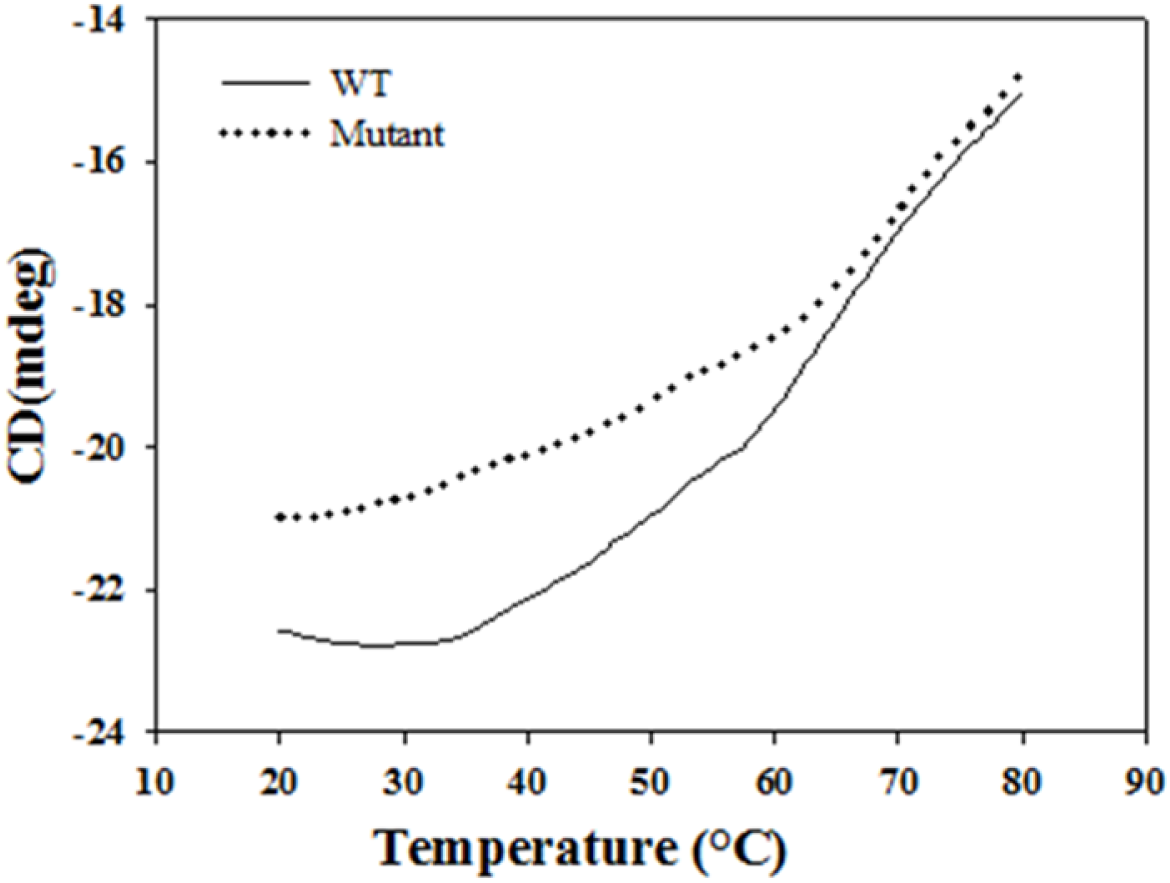
Effect of temperature (with 100 mM of NaCl) on stability of protein.

**Figure S3:**
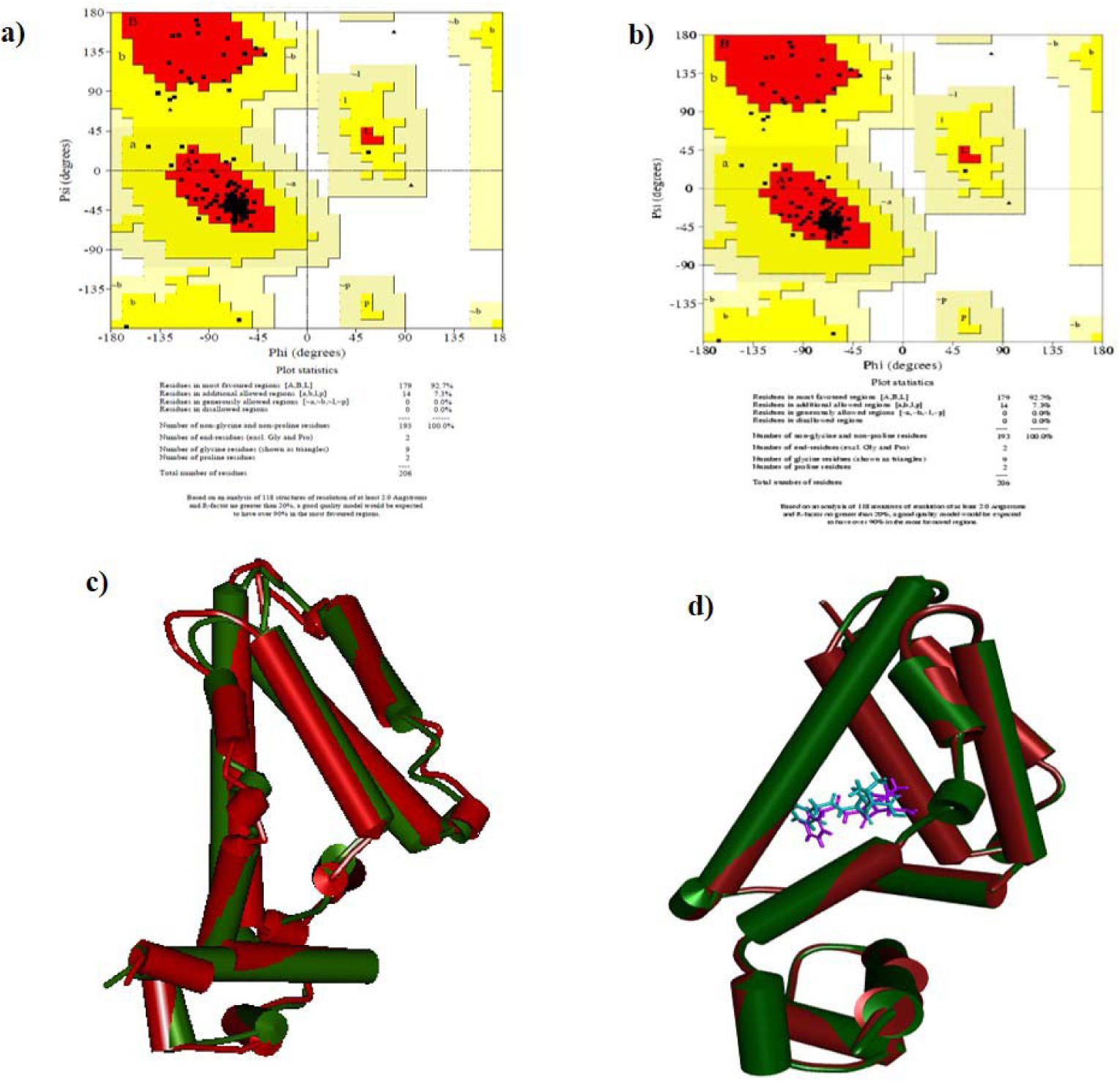
The Ramachandran plots depicting that the modeled structures (MtrR) are stable as most of the amino acids fall in the permissible zone. a) WT, b) H105Y, c) Superimposed structure of WT (Red) and H105Y mutant MtrR (Green), and d) H105Y mutant docked with penicillin (green) superimposed on wild type MtrR docked with penicillin (red).

**Figure S4:**
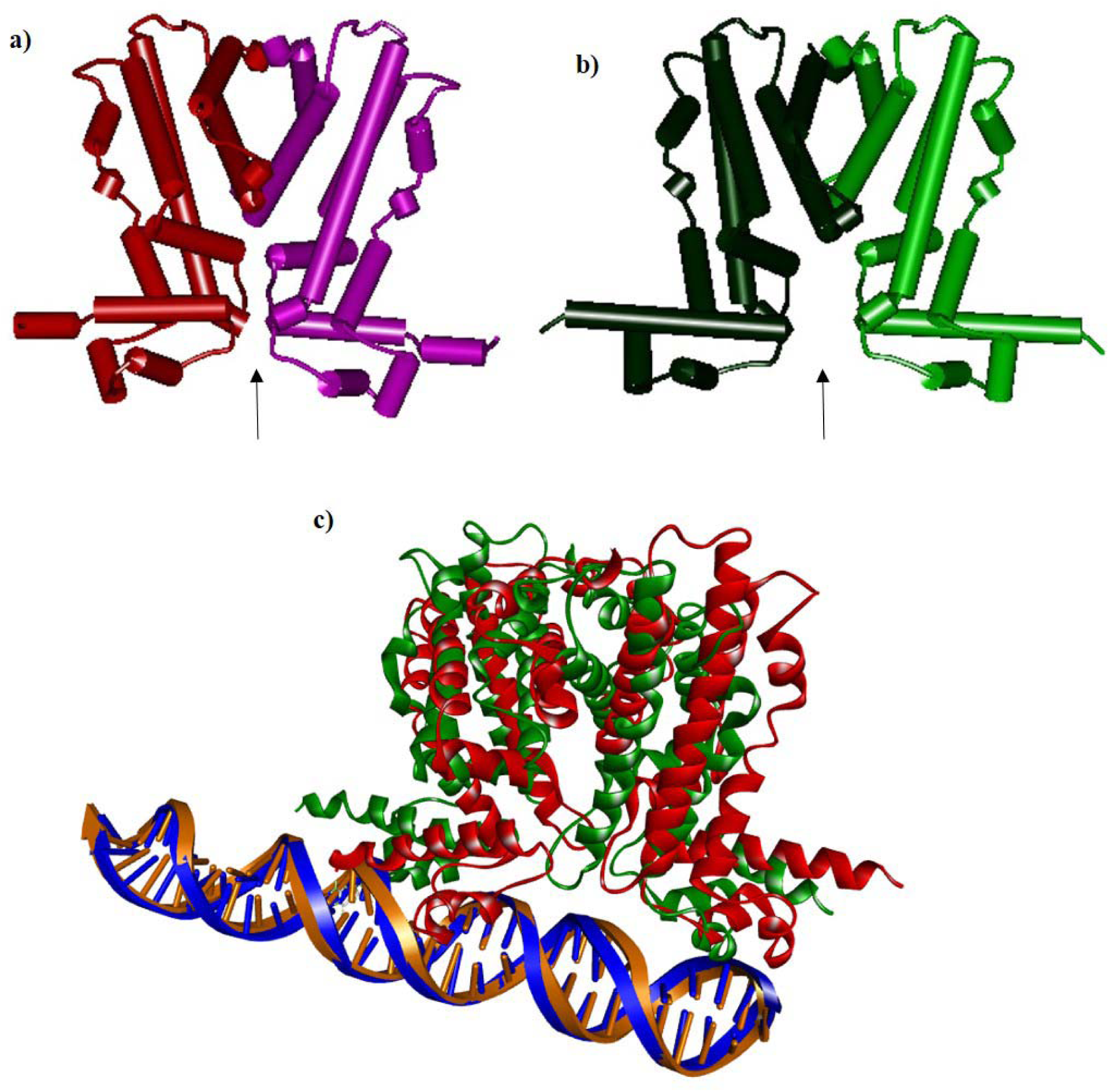
Structure of dimer of WT (a) and mutant (b) showing change in center to center distance between two monomeric units; Superimposed structure of mutant (green) docked with DNA (brown) on WT Red) docked with DNA (blue) (c).

